## Supplemental Figures for "Engineered Immunogens to Elicit Antibodies Against Conserved Coronavirus Epitopes"

### Supplementary Online Materials

**Supplementary Table 1. Design characteristics and initial experimental screening data for all the epitope scaffolds tested experimentally.** FP, S2hlx and S2hlx-Ex epitope scaffolds originate from diverse parent proteins. Epitope scaffolds were expressed recombinantly in *E. Coli* and tested for binding to spike$^{815\text{-}823}$ and stem helix targeting mAbs by ELISA (+++ denotes high affinity; ++ moderate affinity; + low affinity). SC= side chain; BB=backbone; AF= AlphaFold2.

| | S2hlx-Ex19 | S2hlx-7-DH1057.1 | FP15-DH1058 |
|---|---|---|---|
| PDB ID | 8F5H | 8F5I | 8FDO |
| *Data collection* | | | |
| Space group | *C*2 | *P*1 | $P2_1$ |
| Cell dimensions | | | |
| a, b, c (Å) | 175.6, 28.7, 65.5 | 56.7, 76.4, 79.0 | 48.4, 50.6, 146.2 |
| α, β, γ (°) | 90.0, 106.7, 90.0 | 71.6, 70.0, 88.3 | 90.0, 96.9, 90.0 |
| Resolution (Å) | 44.55-2.30 (2.38-2.30) | 45.68-1.90 (1.97-1.90) | 47.33-2.20 (2.28-2.20) |
| $R_{merge}$ | 0.705 (1.651) | 0.664 (0.933) | 0.745 (1.051) |
| I/σI | 16.1 (2.4) | 29.1 (3.2) | 23.6 (2.3) |
| CC1/2 | 0.765 (0.613) | 0.474 (0.207) | 0.501 (0.529) |
| Completeness (%) | 95.1 (92.6) | 93.9 (87.6) | 98.2 (98.6) |
| Redundancy | 13.5 (14.0) | 4.7 (3.7) | 7.5 (7.5) |
| *Refinement* | | | |
| $R_{work}/R_{free}$ (%) | 25.4/27.5 | 21.1/24.5 | 18.9/22.9 |
| No. atoms | | | |
| Protein | 2,212 | 8,003 | 4,264 |
| Glycan | 0 | 111 | 0 |
| Water | 69 | 1,353 | 189 |
| Average B-factors | | | |
| Proteins | 45.6 | 24.7 | 61.6 |
| Ligands | 46.4 | 44 | 49.5 |
| R.m.s deviations | | | |
| Bond lengths (Å) | 0.007 | 0.008 | 0.006 |
| Bond angles (Å) | 1.00 | 0.86 | 0.85 |
| Ramachandran | | | |
| Favored (%) | 99.2 | 98.5 | 97.1 |
| Allowed (%) | 0.8 | 1.5 | 2.7 |
| Outliers (%) | 0 | 0 | 0.2 |

**Supplementary Table 2. Crystallographic table.**

| | Patient # | Vaccine | Days post vaccination | COVID strain | Variant of concern | Days post symptom onset |
|---|---|---|---|---|---|---|
| Vaccinated | 1 | Ad26.Cov2.S plus BNT162b2 | 42 | | | |
| | 2 | mRNA-1273 | 28 | | | |
| | 3 | mRNA-1273 | 29 | | | |
| | 4 | mRNA-1273 | 20 | | | |
| | 5 | mRNA-1273 | 14 | | | |
| | 6 | mRNA-1273 | 44 | | | |
| | 7 | mRNA-1273 | 13 | | | |
| | 8 | BNT162b2 | 52 | | | |
| | 9 | BNT162b2 | 33 | | | |
| | 10 | BNT162b2 | 26 | | | |
| | 11 | BNT162b2 | 26 | | | |
| | 12 | BNT162b2 | 33 | | | |
| Infected | 13 | | | B.1.2 | | 21 |
| | 14 | | | B.1.2 | | 28 |
| | 15 | | | AY.103 | DELTA | 14 |
| | 16 | | | AY.118 | DELTA | 21 |
| | 17 | | | AY.44 | DELTA | 21 |
| | 18 | | | AY.118 | DELTA | 28 |
| | 19 | | | AY.44 | DELTA | 28 |
| | 20 | | | BA.1.1 | OMICRON | 21 |
| | 21 | | | BA.1.1 | OMICRON | 21 |
| | 22 | | | BA.1.1 | OMICRON | 28 |
| | 23 | | | BA.1.1 | OMICRON | 28 |
| | 24 | | | BA.1.1 | OMICRON | 7 |
| Vaccinated + Infected | 25 | Ad26.Cov2.S | 51 | AY.103 | DELTA | 14 |
| | 26 | Ad26.Cov2.S | 192 | AY.44 | DELTA | 21 |
| | 27 | Ad26.Cov2.S | 137 | AY.103 | DELTA | 21 |
| | 28 | Ad26.Cov2.S | 96 | AY.44 | DELTA | 28 |
| | 29 | Ad26.Cov2.S | 193 | BA.1.1 | OMICRON | 28 |
| | 30 | mRNA-1273 | 376 | BA.1.1 | OMICRON | 21 |
| | 31 | BNT162b2 | 65 | B.1.1.207 | | 28 |
| | 32 | BNT162b2 | 38 | AY.44 | DELTA | 21 |
| | 33 | BNT162b2 | 131 | AY.44 | DELTA | 28 |
| | 34 | BNT162b2 | 41 | AY.44 | DELTA | 28 |
| | 35 | BNT162b2 | 516 | BA.2.12.1 | OMICRON | 28 |
| | 36 | BNT162b2 | 437 | BA.2.12.1 | OMICRON | 28 |

**Supplementary Table 3. SARS-CoV-2 strain and vaccine information of patient samples analyzed for binding to native protein domains and the engineered epitope scaffolds.**

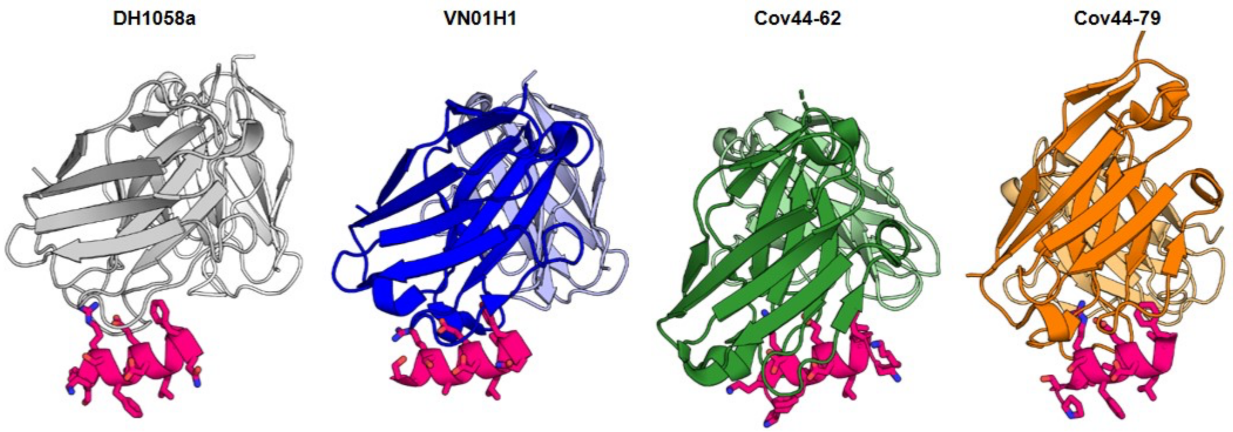


**Supplementary Figure 1. Antibodies with broad recognition against diverse coronaviruses target the conserved spike$^{815-823}$ epitope of SARS-CoV-2.** Known structures of DH1058 (*grey*), VN01H1 (*blue*), Cov44-62 (*green*), and Cov44-79 (*orange*) mAbs bound to their spike epitope (*magenta*). Epitopes are centered on residues 813-824 with key contacts made with virus residues R815, E819 and F823.

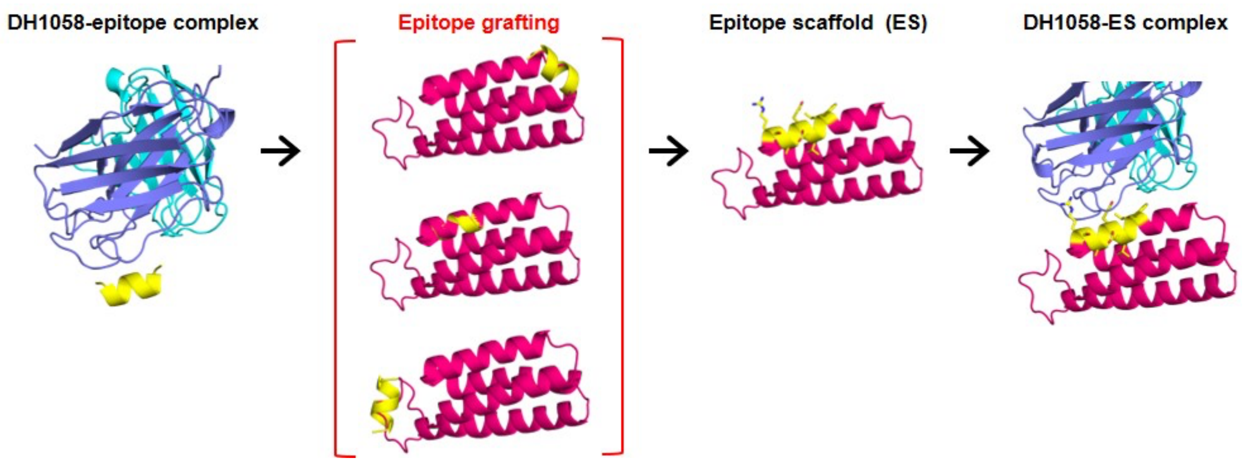


**Supplementary Figure 2. Workflow for computational design of spike$^{815-823}$ epitope scaffolds by side chain grafting.** Using the crystal structure of antibody DH1058 (*light purple* and *cyan*) bound to its epitope (*yellow*), candidate scaffolds (*magenta*) were queried computationally to identify proteins with exposed backbone regions that closely matched (<0.5Å RMS) the structure of the antibody-bound epitope. On proteins that have regions with high structural mimicry to the DH1058-bound epitope, the epitope sequence (*yellow*) replaced the native scaffold to generate an epitope scaffold (ES); additional mutations were introduced into the scaffold to accommodate the grafted epitope, to prevent clashes with the DH1058 mAb in the modeled antibody-ES complex, and to revert any unnecessary scaffold mutations introduced at the automated computational design stage.

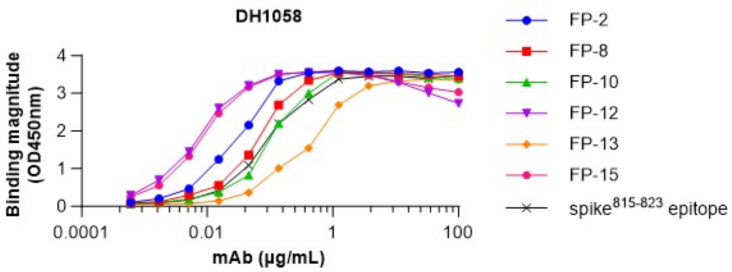


**Supplementary Figure 3. Initial binding screen of designed spike$^{815\text{-}823}$ epitope scaffolds to DH1058 mAb.** ELISA binding of DH1058 mAb to immobilized epitope scaffolds and a synthetic peptide encompassing the spike$^{815\text{-}823}$ epitope (SARS-CoV-2 spike residues 808-833).

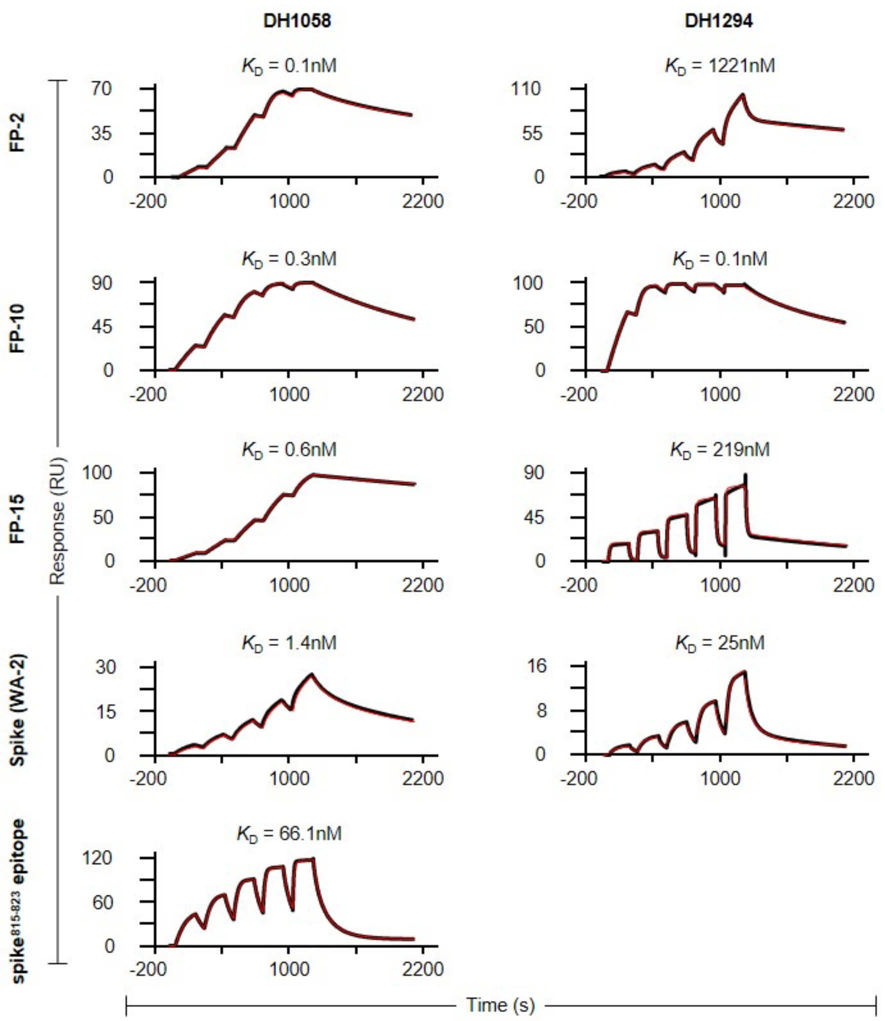


**Supplementary Figure 4. Surface Plasmon Resonance binding curves to determine the dissociation constants of spike$^{815\text{-}823}$ epitope scaffolds for DH1058 and DH1294 mAbs.** Acquired data is shown in *black* and the curve fit is in *red*. Data is representative of at least two independent experiments.

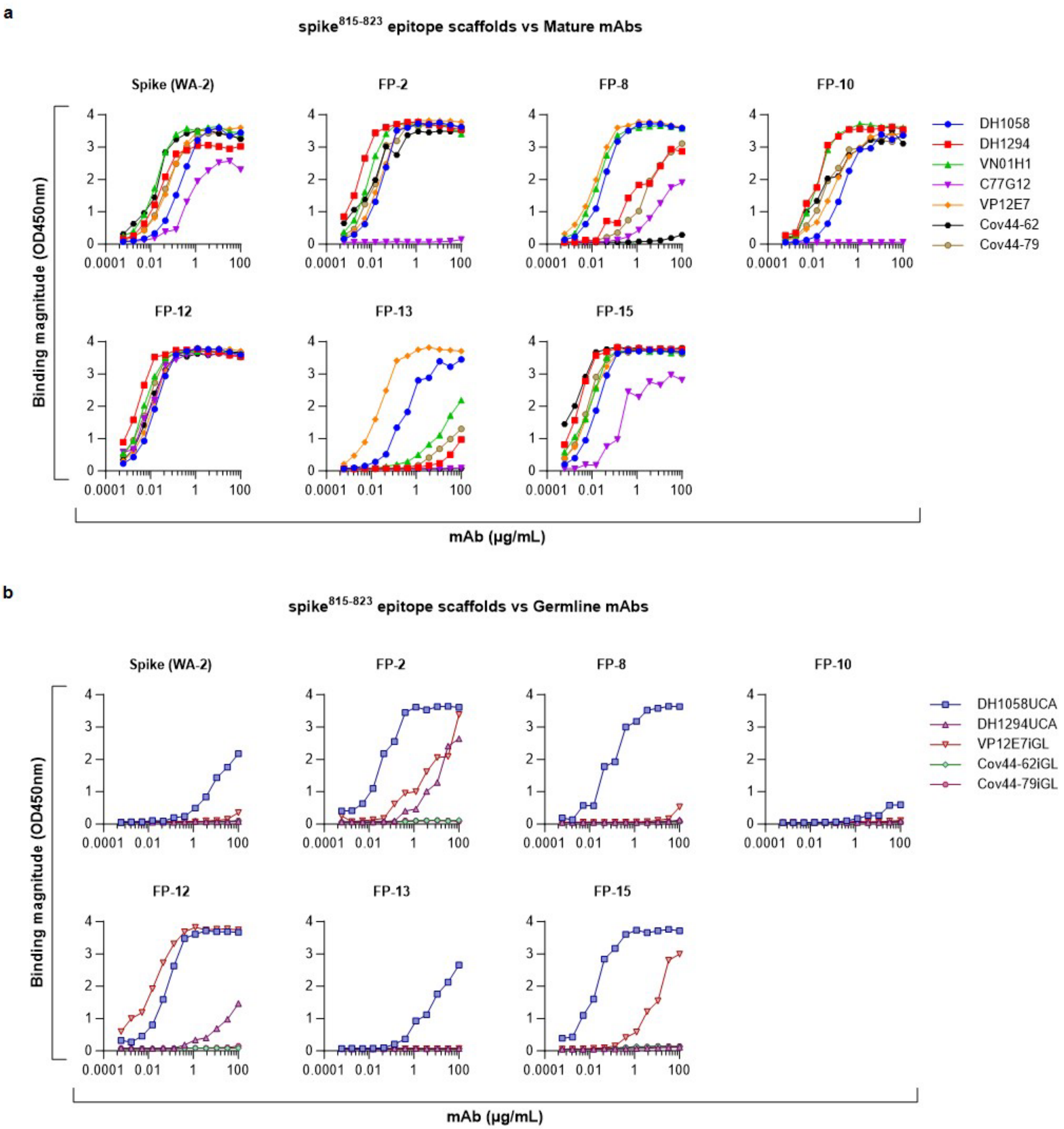


**Supplementary Figure 5. Epitope scaffolds bind to diverse mature and germline antibodies that target the spike$^{815\text{-}823}$ epitope. (a)** ELISA binding of mAbs DH1058, DH1294, VN01H1, C77G12, VP12E7, Cov44-62, and Cov44-79 to FP ESs and to a synthetic peptide encompassing the spike$^{815\text{-}823}$ epitope (SARS-CoV-2 spike residues 808-833). **(b)** ELISA binding of mAbs DH1058 UCA, DH1294 UCA, VP12E7 iGL, Cov44-62 iGL, and Cov44-79 iGL to FP ESs and to a synthetic peptide encompassing the spike$^{815-823}$ epitope (SARS-CoV-2 spike residues 808-833).

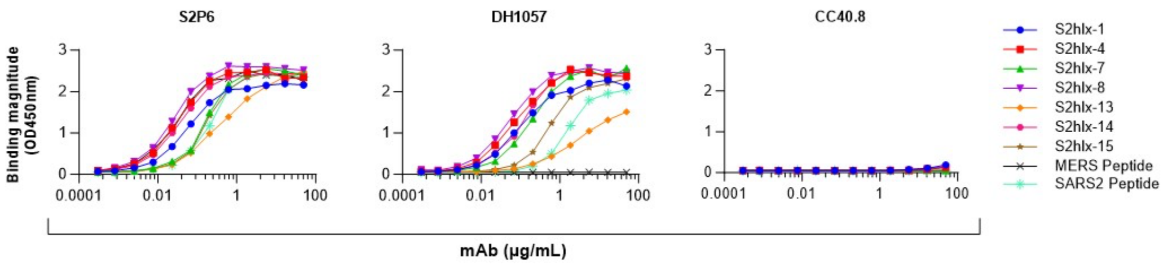


**Supplementary Figure 6. S2hlx ESs bind to S2P6 and DH1057.1 mAbs, but not to CC40.8 mAb.** ELISA binding of mAbs S2P6, DH1057.1 and CC40.8 to S2hlx ESs measured by ELISA. Binding was compared to that of synthetics stem helix peptides derived from SARS-CoV2 ($^{1147}$SFKEELDKYFKNHTS$^{1161}$) and MERS ($^{1230}$DFQDELDEFFKNVST$^{1244}$). The synthetic peptide did not contain residue L1145 (SARS-CoV-2 numbering) that is critical for epitope binding by CC40.8.

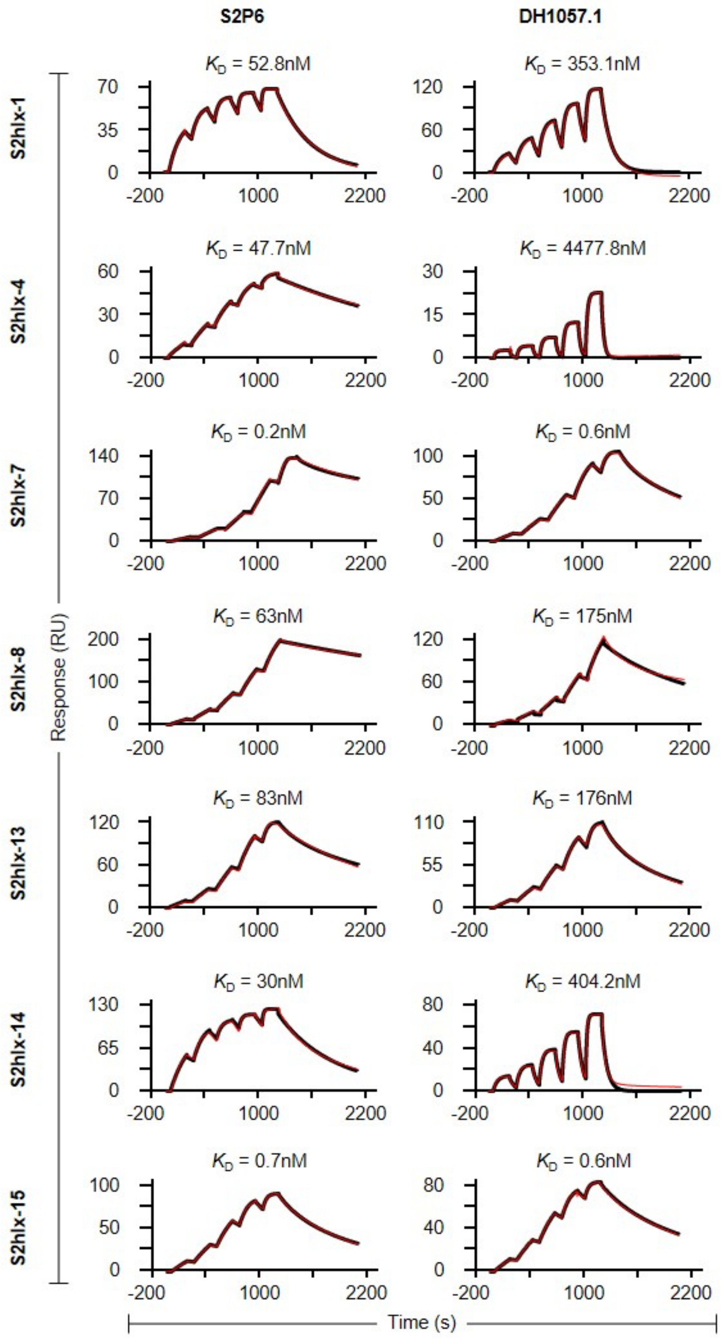


**Supplementary Figure 7. Surface Plasmon Resonance curves of S2hlx epitope scaffolds binding to S2P6, S2P6iGL, DH1057.1 and DH1057UCA mAbs.** Acquired data is shown in *black* and the curve fit is in *red*.

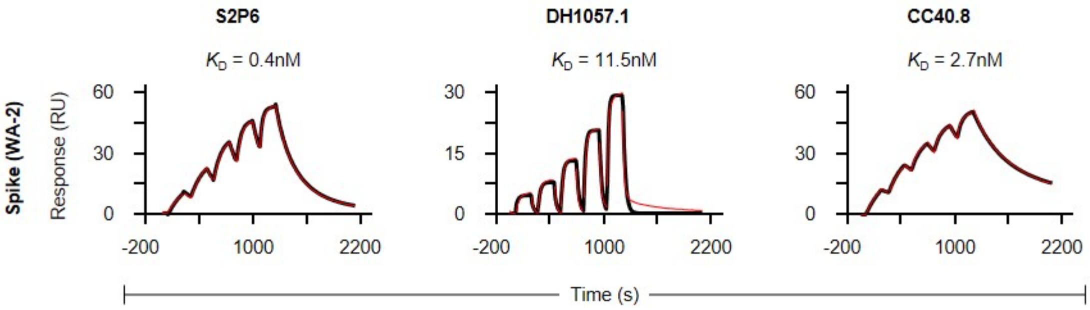


**Supplementary Figure 8. Surface Plasmon Resonance curves of Spike (WA-2) proteins binding to S2P6, DH1057.1 and CC40.8 mAbs.** Acquired data is shown in *black* and the curve fit is in *red*.

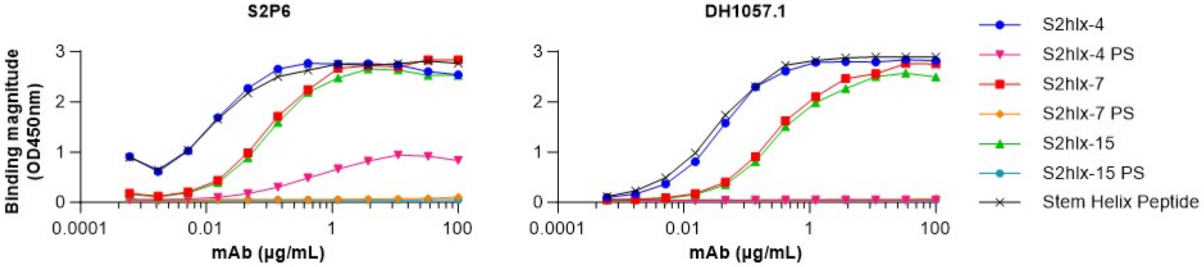


**Supplementary Figure 9. Binding of S2hlx epitope scaffolds to target antibodies is mediated by the grafted epitope residues.** ELISA binding of S2P6 and DH1057 mAbs to S2hlx epitope scaffolds and to the parent scaffolds (PS) that served as templates for epitope grafting.

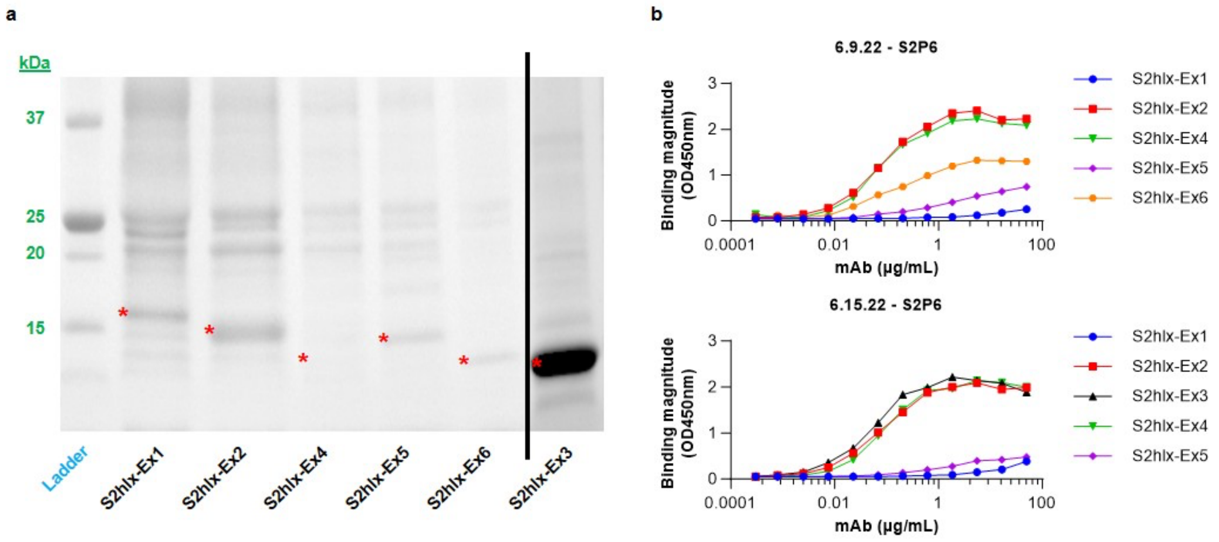


**Supplementary Figure 10. Recombinant expression and S2P6 mAb binding of S2hlx-Ex epitope scaffolds designs generated by backbone grafting. (a)** SDS-PAGE gel showed limited expression of initial S2hlx-Ex epitope scaffolds. **(b)** Binding of mAb S2P6 to immobilized S2hlx-Ex epitope scaffolds measured by ELISA.

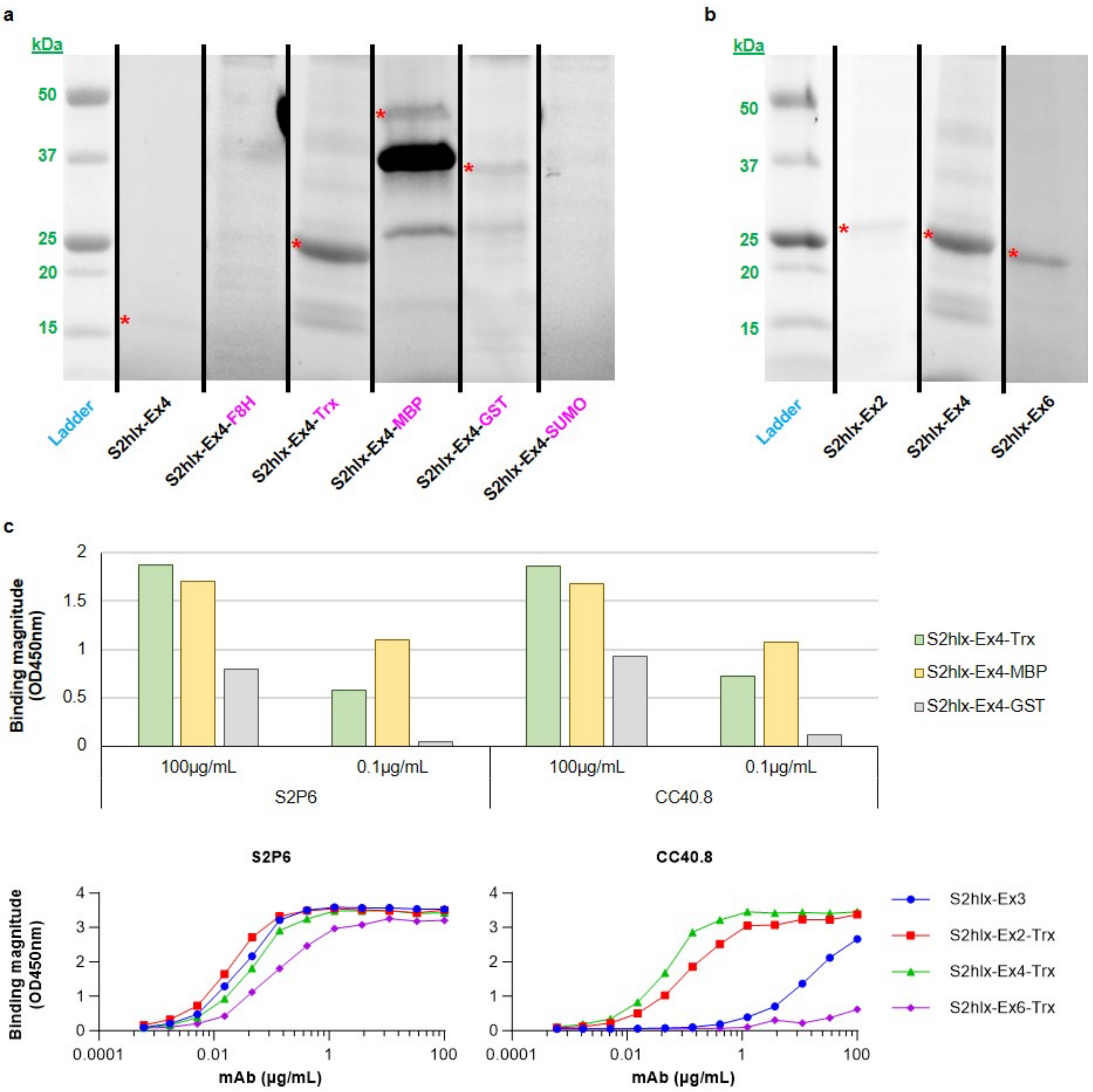


**Supplementary Figure 11. Tagged S2hlx-Ex epitope scaffolds showed improved expression and antibody binding. (a)** SDS-PAGE gel showing the expression of the initial S2hlx-Ex4 design and of different versions fused to the tags highlighted in *magenta*. **(b)** SDS-PAGE gel showing expression of Trx-tagged versions of S2hlx-Ex2, Ex4, and Ex6. * marks expected molecular weight bands. **(c)** Binding of S2P6 and CC40.8 mAbs to tagged S2hlx-Ex epitope scaffolds measured by ELISA.

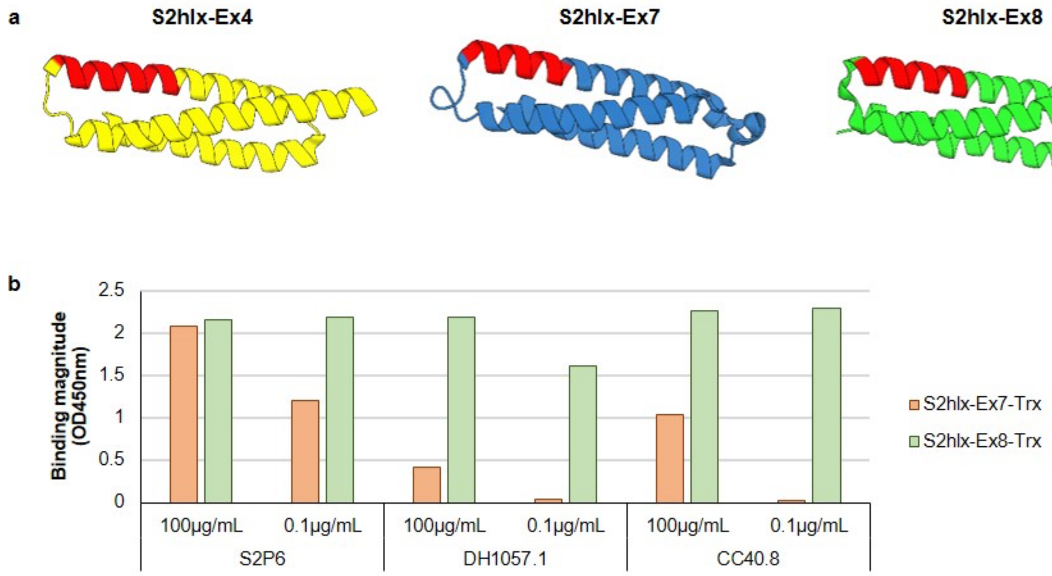


**Supplementary Figure 12. Binding of epitope scaffolds designed on structural homologs of the S2hlx-Ex4 parent scaffold. (a)** Structures of the S2hlx-Ex4 epitope scaffold and of two related designs, S2hlx-Ex7 and S2hlx-Ex8, based on structurally homologous parent scaffolds (PDBids: 2qyw, 1lvf, and 3onj). The grafted epitope is shown in *red*. **(b)** ELISA binding of antibodies S2P6, DH1057, and CC40.8 to tagged epitope scaffolds from **(a)**.

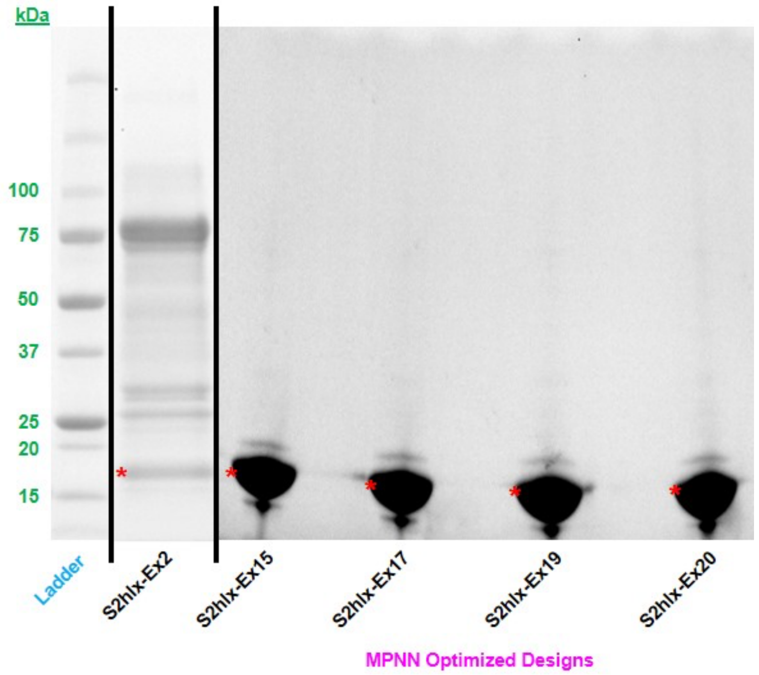


**Supplementary Figure 13. Expression of S2hlx-Ex2 and related successful MPNN designs.** SDS-PAGE gel showing the expression of the initial S2hlx-Ex2 design and of different MPNN versions.

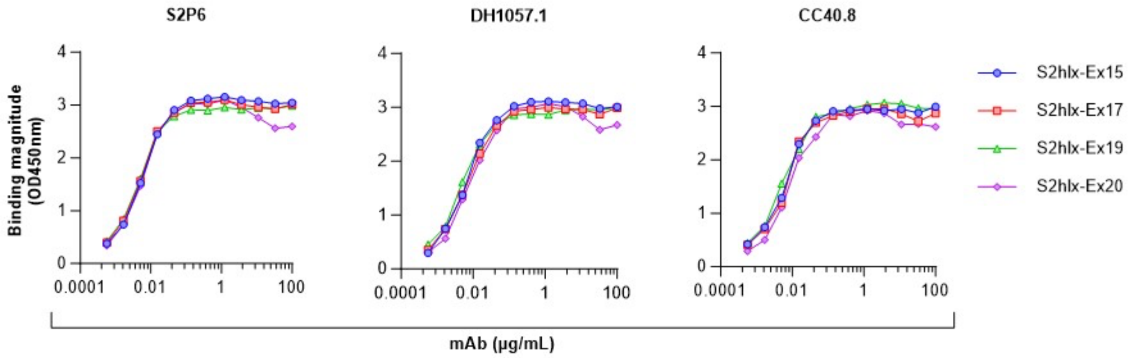


**Supplementary Figure 14. S2hlx-Ex2 based MPNN designs show broad binding.** Binding of S2P6, DH1057.1 and CC40.8 mAbs to MPNN designs based on S2hlx-Ex2 epitope scaffolds measured by ELISA.

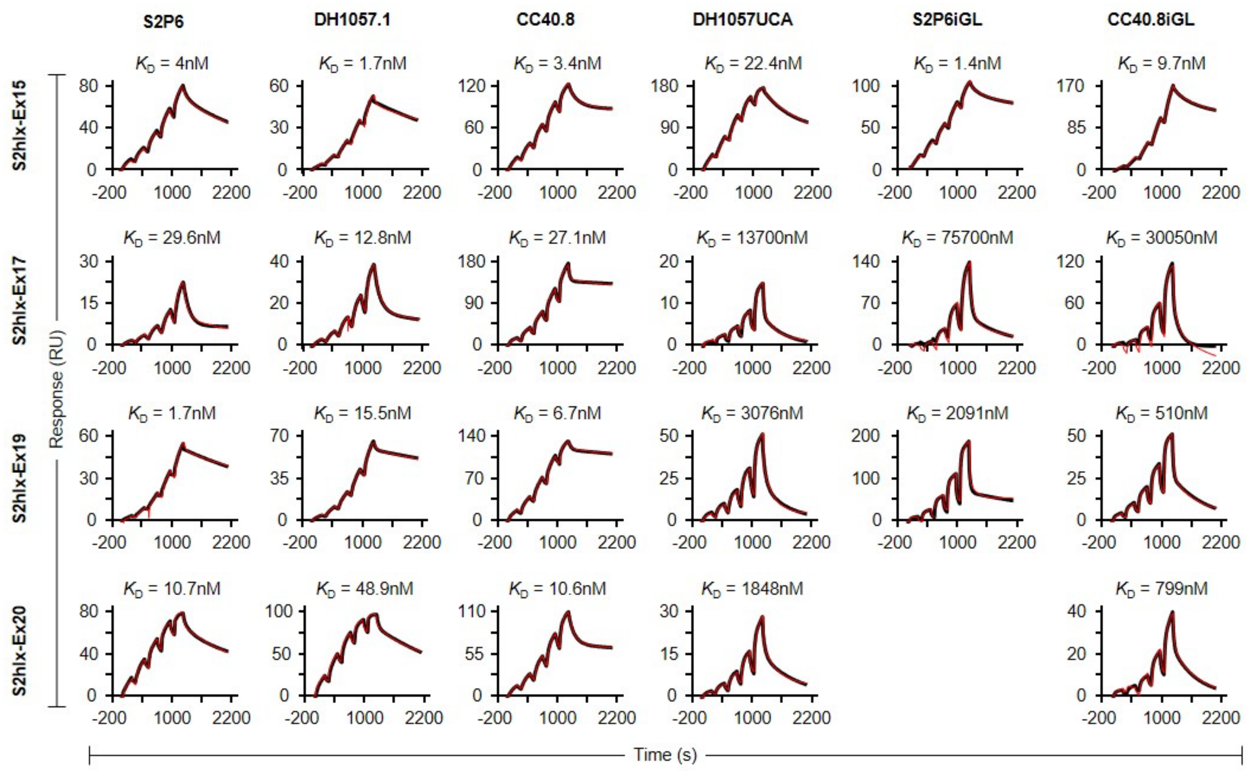


**Supplementary Figure 15. Surface Plasmon Resonance curves of S2hlx-Ex2 derived epitope scaffolds binding to S2P6, S2P6iGL, DH1057.1, DH1057 UCA, CC40.8 and CC40.8iGL antibodies.** Binding kinetics of S2hlx-Ex2 derived epitope scaffolds to S2P6, DH1057, and CC40.8 mAbs and to their germline variants as measured by Surface Plasmon Resonance (SPR).

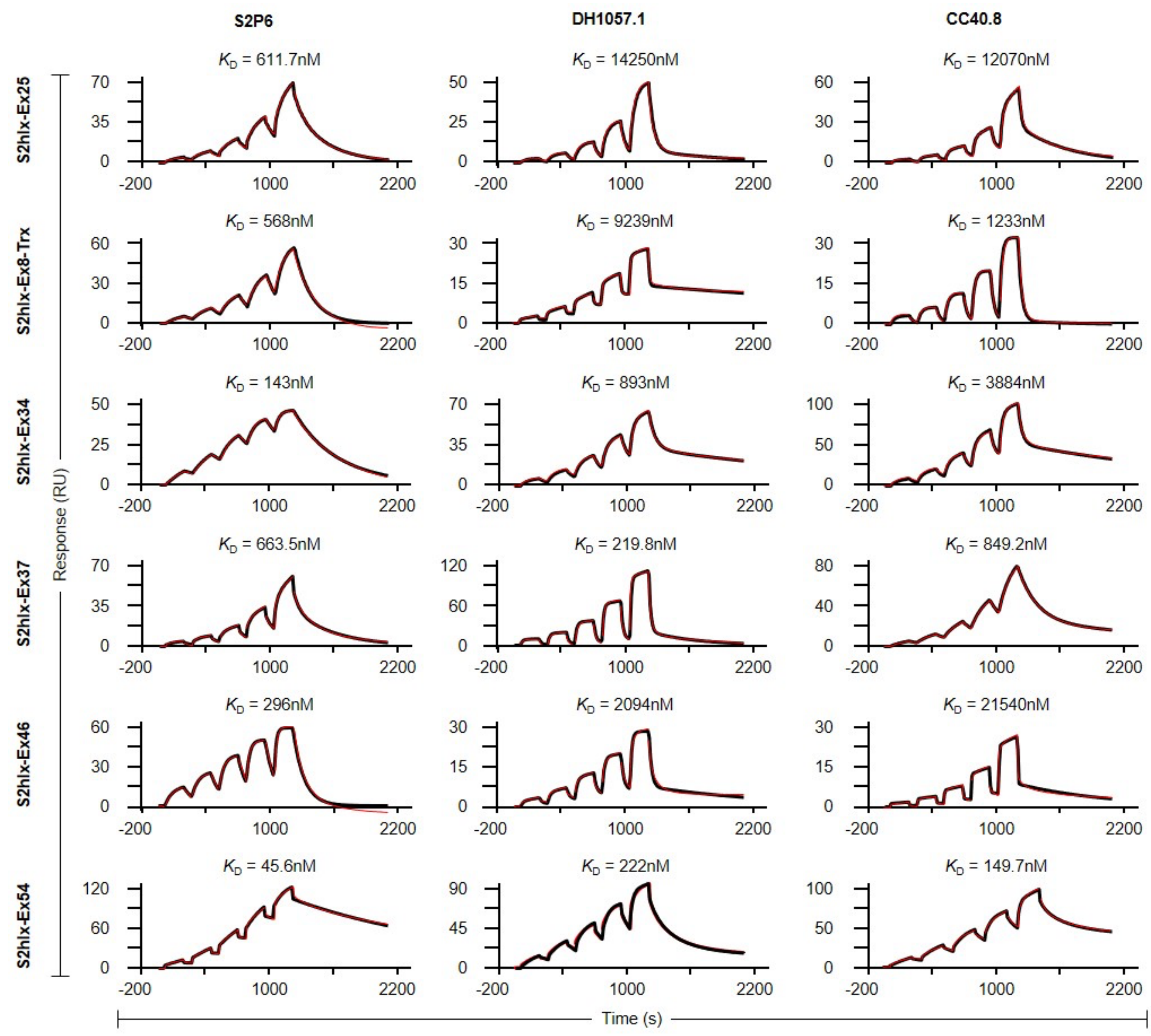


**Supplementary Figure 16. Binding affinities of S2hlx-Ex4, Ex6, and Ex3 derived epitope scaffolds to S2P6, DH1057, and CC40.8 mAbs.** Binding kinetics of S2hlx-Ex4, Ex6, and Ex3 derived epitope scaffolds to S2P6, DH1057, and CC40.8 as measured by Surface Plasmon Resonance (SPR).

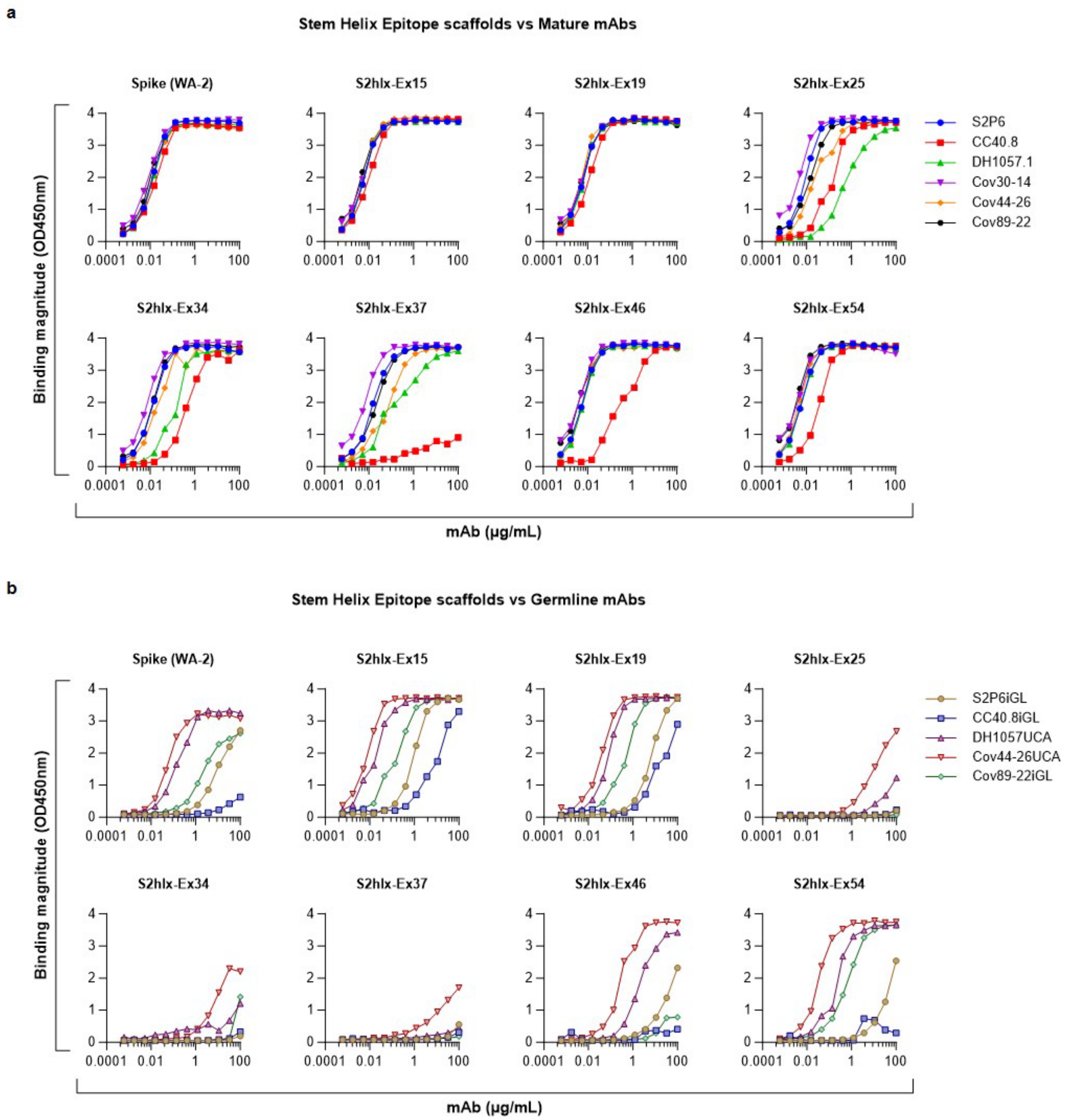


**Supplementary Figure 17. Epitope scaffolds bind to diverse mature and germline antibodies that target the stem helix epitope. (a)** ELISA binding of mAbs S2P6, DH1057.1, CC40.8, Cov30-14, Cov44-26, and Cov89-22 to S2hlx-Ex ESs and to Spike (WA-2) protein. **(b)** ELISA binding of mAbs S2P6iGL, DH1057 UCA, CC40.8 iGL, Cov44-26 UCA, and Cov89-22 iGL to S2hlx-Ex ESs and to Spike (WA-2) protein.

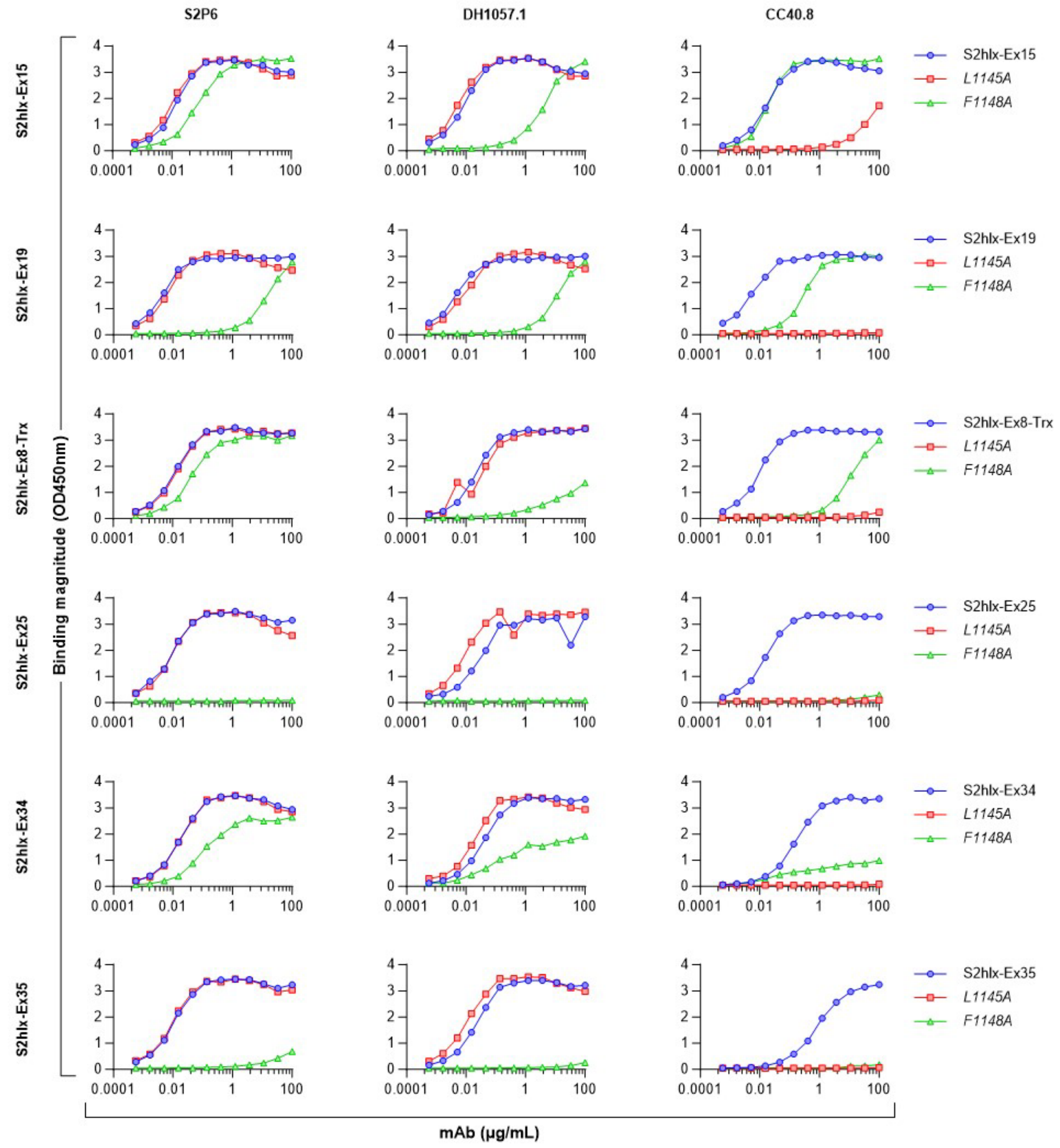
S2P6
DH1057.1
CC40.8
S2hlx-Ex15
S2hlx-Ex19
S2hlx-Ex8-Trx
S2hlx-Ex25
S2hlx-Ex34
S2hlx-Ex35
Binding magnitude (OD450nm)
mAb (µg/mL)
0.0001
0.01
1
100
L1145A
F1148A

**Supplementary Figure 18. Binding specificity of S2hlx-Ex epitope scaffolds for representative epitopes against the stem helix epitope.** Indicated spike mutations known to affect the binding of stem helix were introduced in the respective epitope scaffolds.

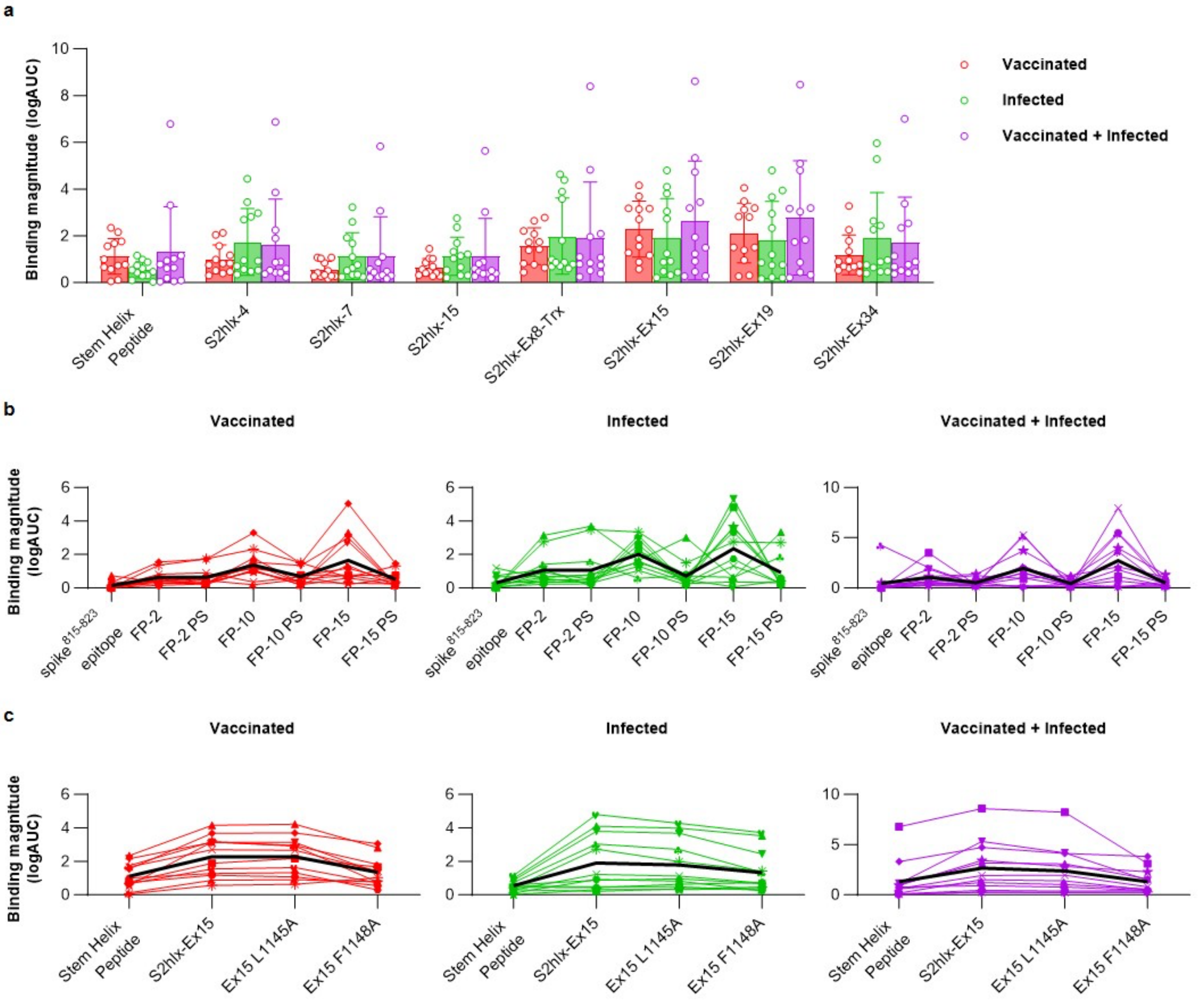


**Supplementary Figure 19. Reactivity of designed epitope scaffolds to patient sera with pre-existing immunity to the SARS-CoV-2 spike. (a)** Binding of stem helix peptide and select S2hlx and S2hlx-EX epitope scaffolds. **(b)** Binding of stem helix peptide, S2hlx-EX-15 and S2hlx-EX-15 epitope mutants that reduce binding to either the CC40.8-class of mAbs (LA) or to both the CC40.8 and S2P6 classes of mAbs (FA). **(c)** Binding to spike$^{815\text{-}823}$ epitope scaffolds and to the parent scaffolds they were derived from and that lack the grafted epitope. Lines represent individual measurements for each sample in the group.

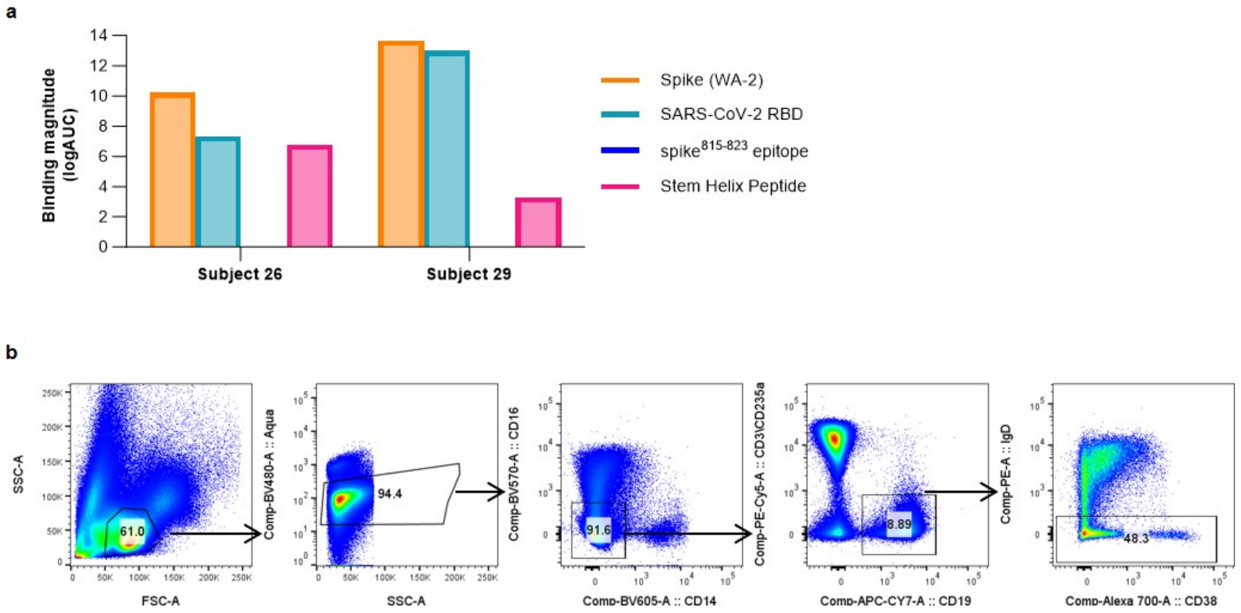


**Supplementary Figure 20. Isolation of stem helix epitope-specific memory B cells from vaccinated then infected subjects with stem helix scaffolds. (a)** ELISA binding of sera from indicated subjects with pre-existing SARS-CoV-2 immunity acquired by vaccination followed by infection to recombinant WA-2 spike and RBD or to synthetic peptides containing the spike$^{815\text{-}823}$ epitope (residues 808-833) or the stem helix (1140-1164). **(b)** Comparison between sera binding to the synthesized stem helix peptide with S2hlx-Ex19 ES, and with versions of this design where the epitope is mutated to reduce binding to either CC40.8 class antibodies (L1145A), to both CC40.8 and S2P6 classes of antibodies (F1148A), or where the epitope is replaced with the sequence from the parent scaffold it replaced (PShlx). Single lines represent measurements for individual samples. **(c)** Flow cytometric gating strategy for the isolation of memory B cells from PBMCs of subjects.

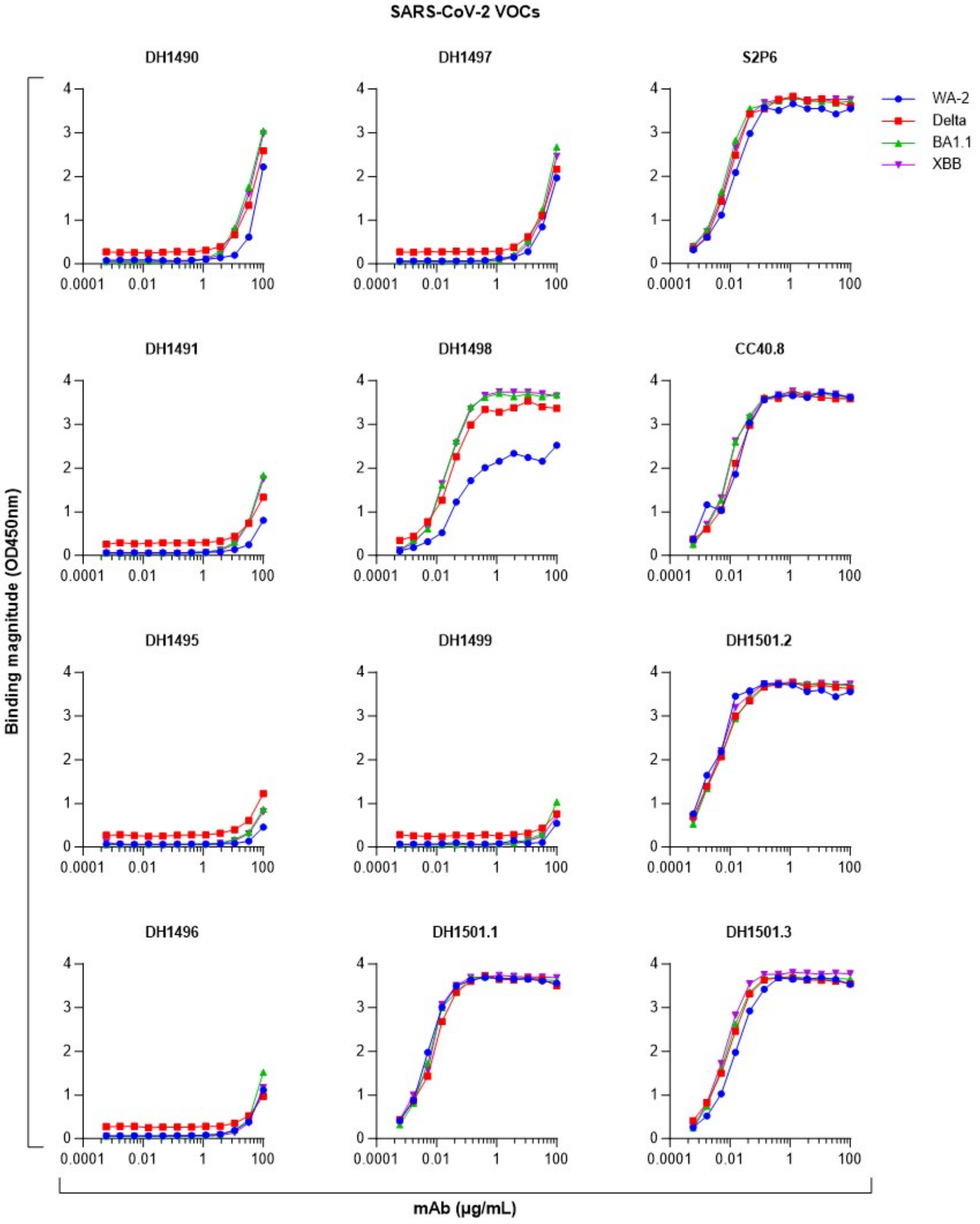


157

158 **Supplementary Figure 21. ELISA binding of isolated stem helix antibodies to SARS-CoV-2**
159 **VOC spikes.**

160

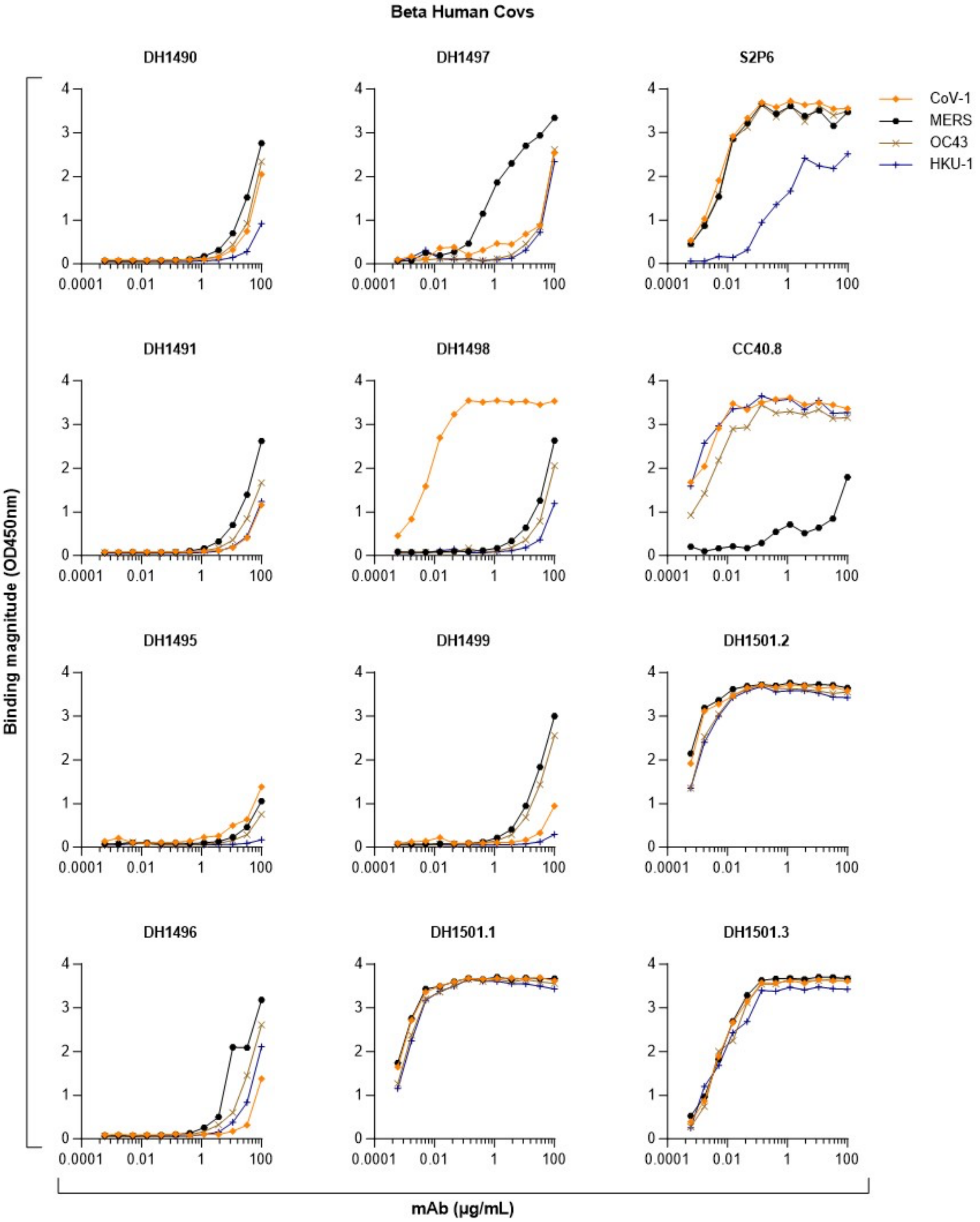


161

162 **Supplementary Figure 22. ELISA binding of isolated stem helix antibodies to human**

163 **betacoronavirus spikes.**

164

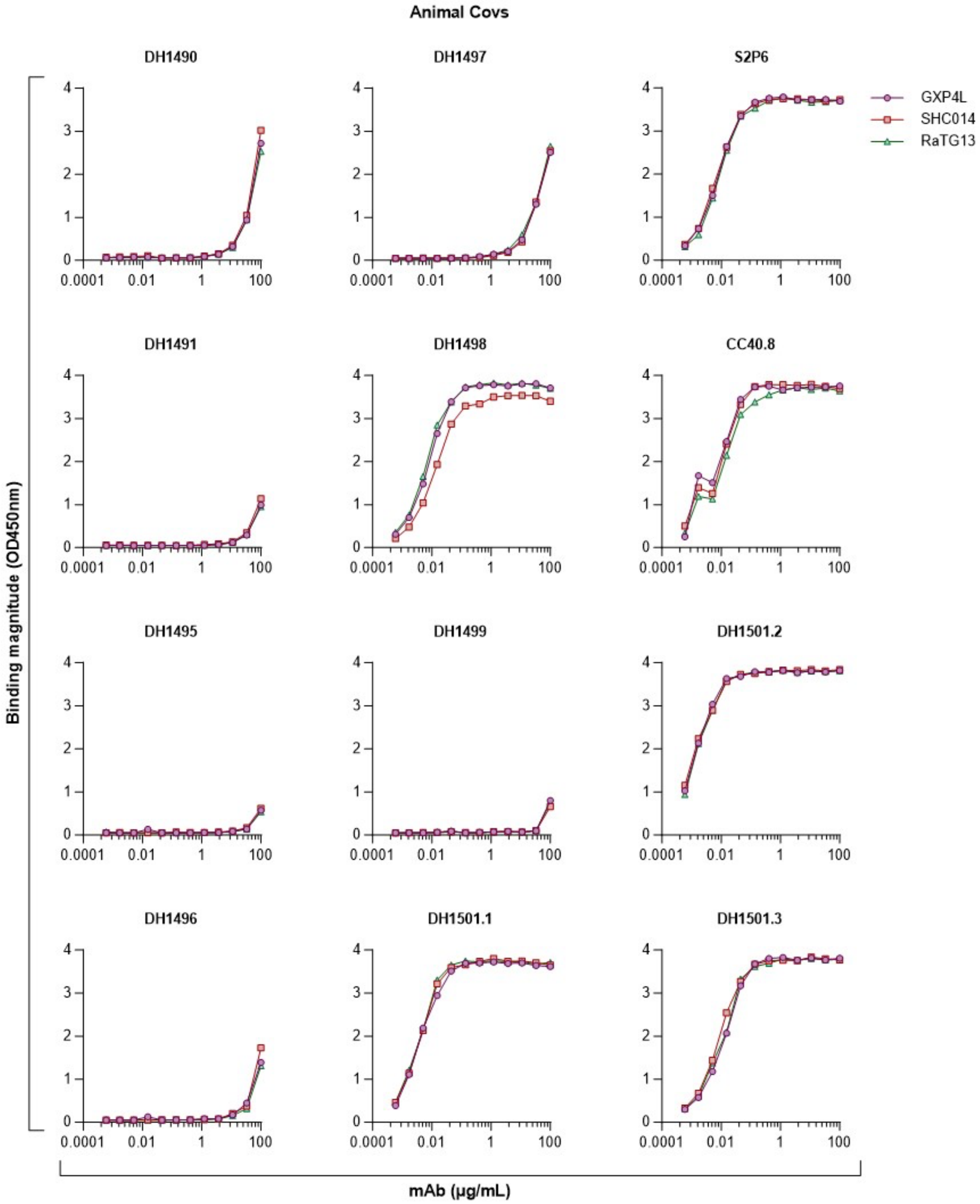


**Supplementary Figure 23. ELISA binding of isolated stem helix antibodies to animal betacoronavirus spikes.**

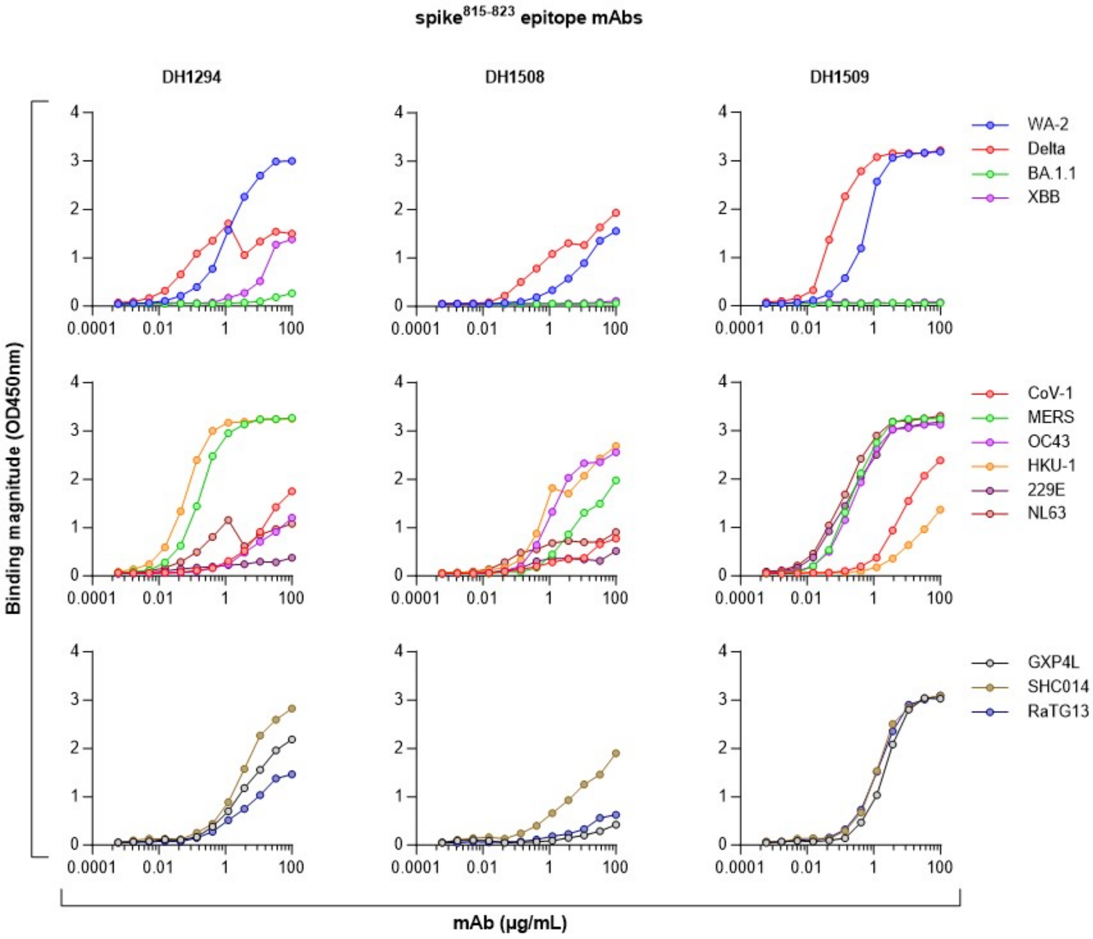


**Supplementary Figure 24. ELISA binding of isolated spike$^{815-823}$ antibodies to human and animal coronavirus spikes.**

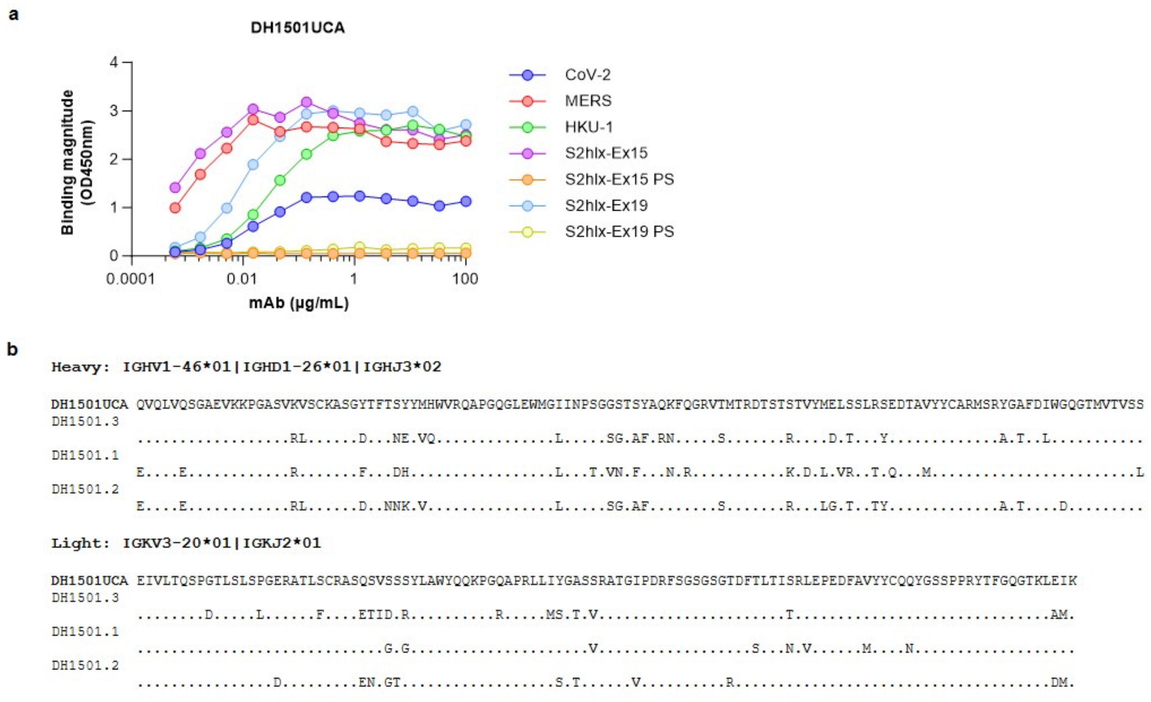


**Supplementary Figure 25. The unmutated common ancestor (UCA) of isolated stem helix antibodies DH1501.1, DH1501.2, and DH1501.3, binds to multiple human coronavirus spikes. (a)** ELISA binding data. **(b)** Sequence alignment between the UCA and the mature antibodies.

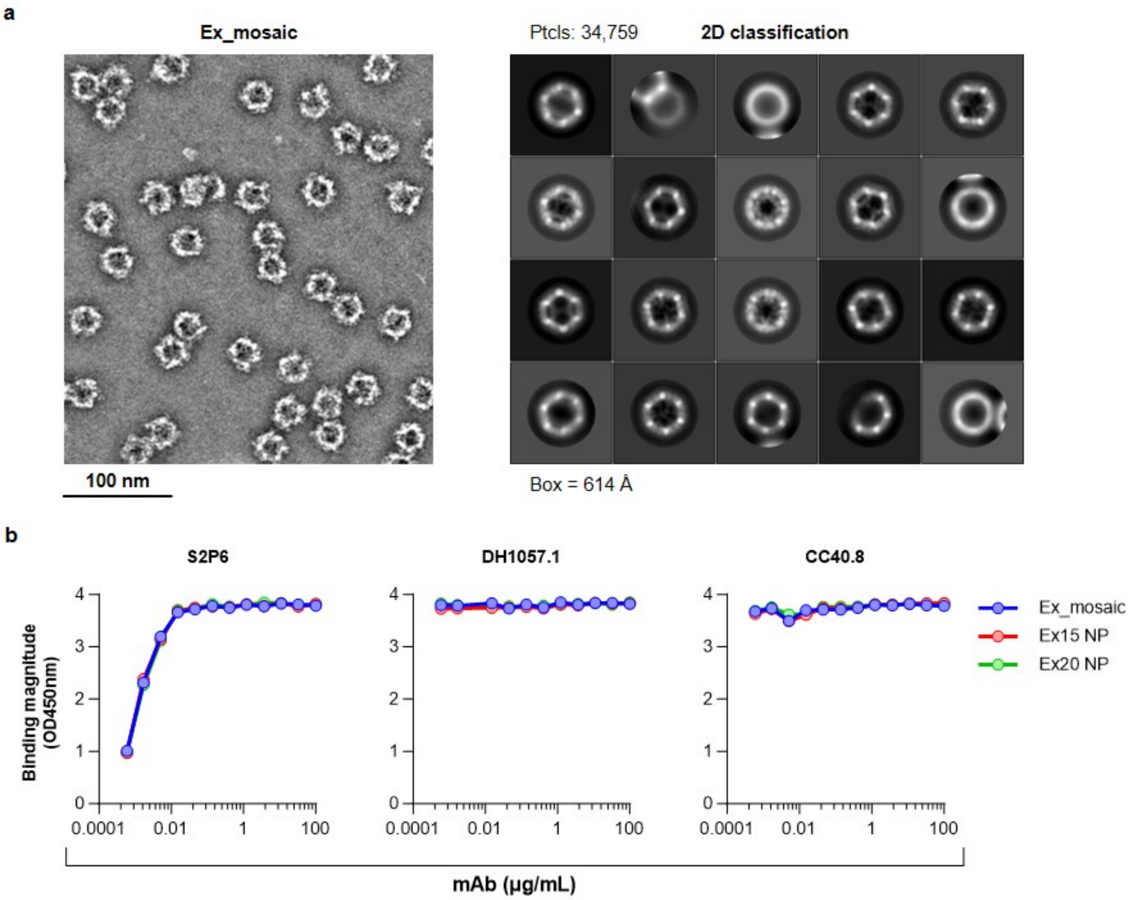


**Supplementary Figure 26. Characterization of epitope scaffold nanoparticles. (a)** NSEM sample image and 2D classification of the Ex_mosaic NP. **(b)** ELISA binding of mi03 nanoparticles displaying the indicated epitope scaffolds to stem helix antibodies S2P6, DH1057.1 and CC40.8.

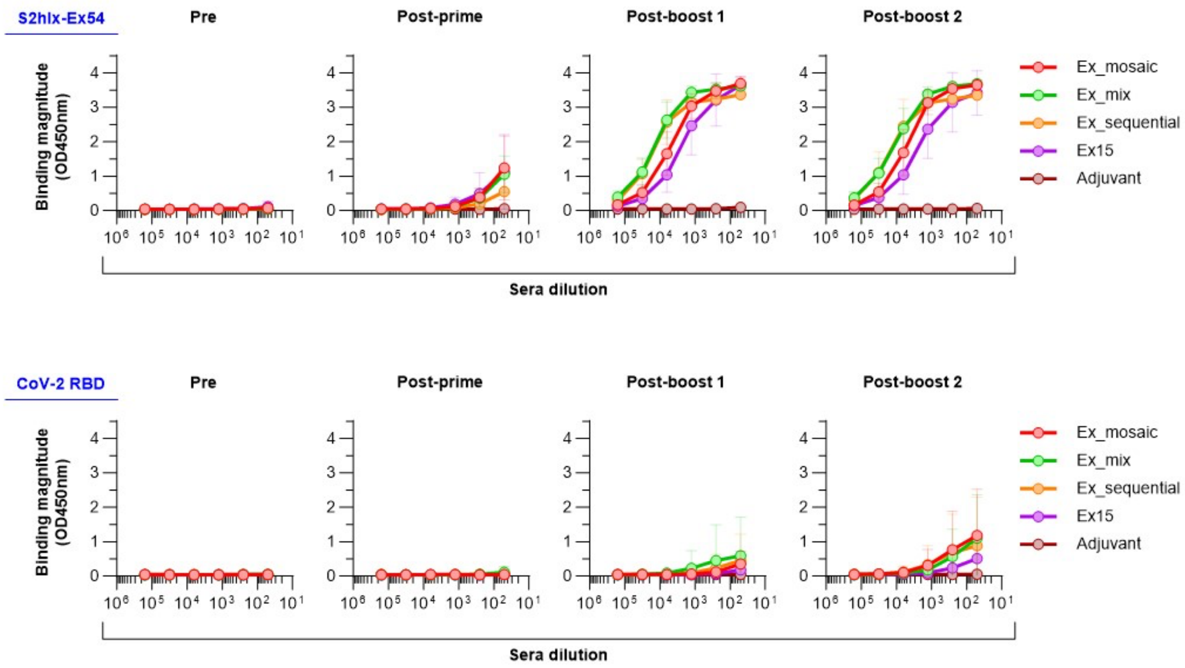


**Supplementary Figure 27. Binding of sera from BALBc immunized mice with stem helix epitope scaffold nanoparticles to an epitope scaffold not used in the immunization (S2hlx-Ex54) and the RBD.**

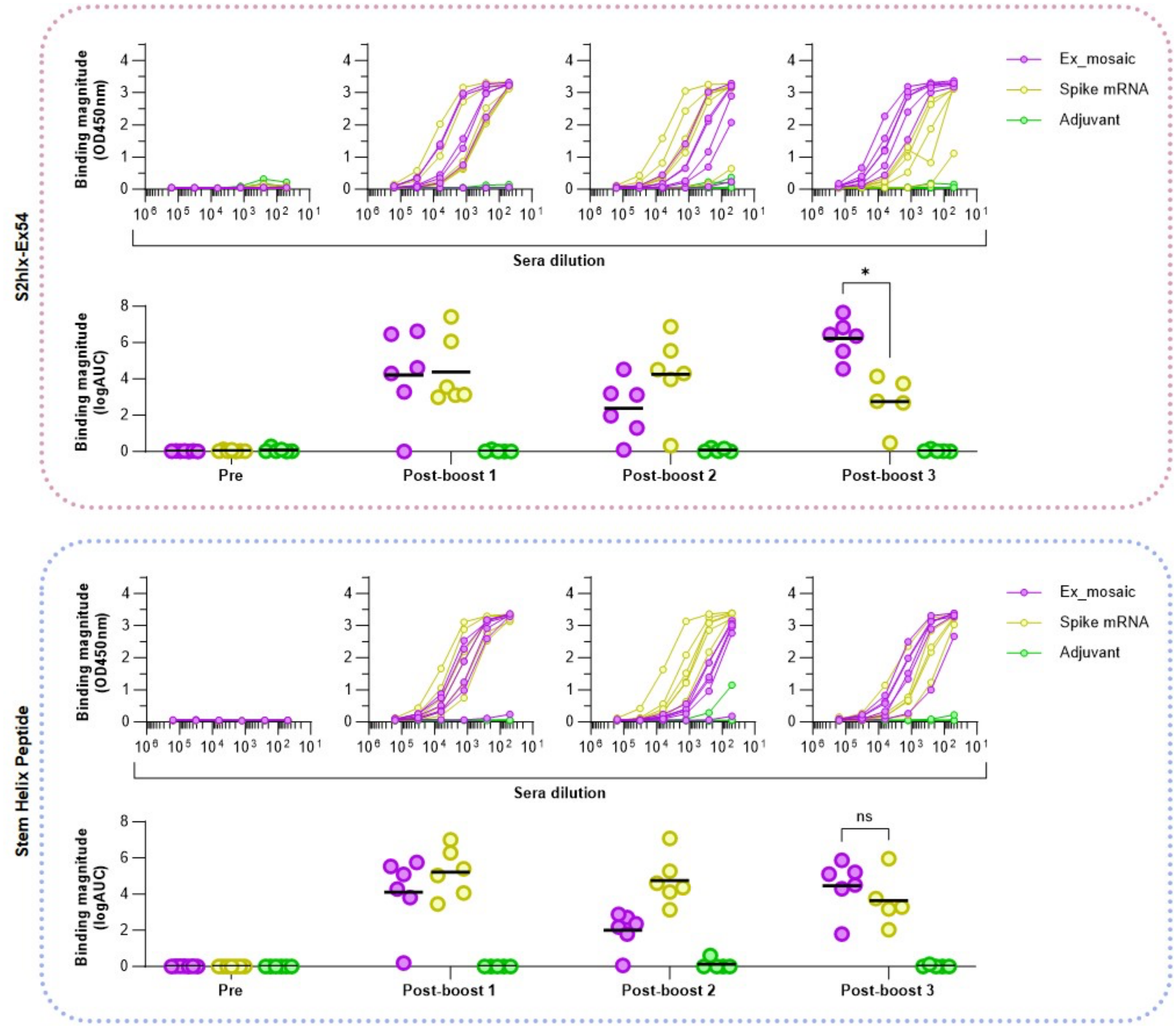


**Supplementary Figure 28. Stem helix epitope specific binding of sera from K18-ACE2 mice primed with spike mRNA and boosted with either the epitope scaffold mosaic NP or spike mRNA.** *Top*: sera binding at indicated time points to a stem helix epitope scaffold not used in the immunizations (S2hlx-Ex54). *Bottom*: sera binding at indicated time points to a synthesized peptide encoding the stem helix epitope.
